# Design and assembly of the 117-kb *Phaeodactylum tricornutum* chloroplast genome

**DOI:** 10.1101/2023.06.05.543740

**Authors:** Emma J.L. Walker, Mark Pampuch, Nelson Chang, Ryan R. Cochrane, Bogumil J. Karas

## Abstract

There is a growing impetus to expand the repository of chassis available to synthetic biologists. The chloroplast genome presents a unique chassis for engineering photosynthetic eukaryotes by virtue of its compact size, lack of epigenetic regulation, and containment within the secluded lipid bilayers of the organelle. The development of the chloroplast as a synthetic biology chassis, however, has been limited by a lack of efficient techniques for whole genome cloning and engineering. Here, we demonstrate two approaches for cloning the 117 kb *Phaeodactylum tricornutum* chloroplast genome that have 90 to 100% efficiency when screening as few as ten *Saccharomyces cerevisiae* colonies following yeast assembly. The first method directly uses PCR-amplified fragments of the genome for yeast assembly, whereas the second method relies upon the pre-cloning of eight overlapping genomic regions into individual plasmids that they can later be released from. The cloned genome can be stably maintained and propagated within *Escherichia coli*, which provides an exciting opportunity for engineering a novel delivery mechanism for bringing DNA directly to the algal chloroplast. As well, one of the cloned genomes was designed to contain a single *Sap*I site within the yeast *URA3* open-reading frame, which can be used to linearize the genome and integrate designer cassettes via golden-gate cloning or further iterations of yeast assembly.

## 1. INTRODUCTION

The twenty-first century is stippled by breakthrough advances within the field of synthetic biology. Over a decade has passed since the first synthetic genome was assembled and transplanted into a cell, bringing to life a novel mycoplasma strain (JCVI-syn1.0; Gibson *et al*., 2010). In the time since, efforts to recode, refactor, minimize, and restructure whole genomes have proven successful at generating microorganisms with unprecedented properties (Chan, Kosuri and Endy, 2005; Endy, 2005; Fredens *et al*., 2019; Dai *et al*., 2020). These accomplishments not only illustrate the plasticity of genomes but also enable scientific advancements that augur well for fundamental research.

Large-scale genome synthesis and transplantation provide the utmost potential for engineering an organism to meet any biologically-possible desire; however, the chassis of such grand transformations have been previously limited to relatively simple or otherwise extensively-studied microorganisms like viruses, bacteria, and yeast (Cello, Paul and Wimmer, 2002; Chan, Kosuri and Endy, 2005; Gibson *et al*., 2010; Annaluru *et al*., 2014; Fredens *et al*., 2019). There is a growing impetus to diversify the repository of chassis available to synthetic biologists that are capable of large-scale genome engineering. Of particular interest is the development of a chassis with photosynthetic capabilities that could serve as a platform for the sustainable generation of bioproducts. Thus far, no such chassis exists with this level of genetic fine-tuning.

Chloroplast genomes could function as unique chassis for engineering photosynthetic eukaryotes. From an assembly perspective, the reduced size of the chloroplast genome (e.g., 110 to 160 kb) and its *nakedness* (i.e., lack of histones; Bock, 2015) makes constructing it from synthetic DNA more feasible than that of most nuclear chromosomes. Several other characteristics make it an industrially-attractive target for encoding an array of bioproducts like medicines, biofuels, bio-fertilizers, and more (Chan *et al*., 2016; Ivleva *et al*., 2016; Hoelscher *et al*., 2018; Li *et al*., 2018; Richter *et al*., 2018; Eseverri *et al*., 2020). In contrast to the nuclear counterpart, the chloroplast genome can be site-specifically modified through innate homologous recombination processes, thereby avoiding any positioning effects, and lacks gene silencing machinery that could otherwise lead to post-insertion transgene repression (Bock, 2015). Furthermore, the chloroplast microenvironment is enveloped by at least two lipid bilayers. This enables the spatial compartmentalization of transgenic pathways, which can increase their efficiency, and the containment of recombinant proteins or metabolic by-products, which could be toxic if present in the cytosol (Huttanus and Feng, 2017). These characteristics enable some chloroplast-encoded transgenes to comprise 2% to 25% of total soluble protein (TSP) levels within photosynthetic plant cells (Meyers *et al*., 2010), though there are records of transgenes exceeding 70% (Oey *et al*., 2009). In comparison, nuclear expression systems often produce 1% to 2% TSP levels within the same systems (Meyers *et al*., 2010).

Development of the chloroplast as a synthetic biology chassis has been limited by a lack of efficient techniques for whole-genome engineering. Past attempts to clone eukaryotic chloroplast genomes have required intensive screening of *Saccharomyces cerevisiae* and/or *Escherichia coli* colonies to identify successfully transformed and intact genomes (Gupta and Hoo, 1991; O’Neill *et al*., 2011). Such cumbersome screening processes are the result of inefficient assembly strategies, which have served as one of the rate-limiting steps in the cloning of chloroplast genomes and thus the creation of designer genomes. To overcome this bottleneck, we sought to design and test several assembly setups in pursuit of a highly efficient and tractable method for cloning the *Phaeodactylum tricornutum* chloroplast genome. *P. tricornutum* has been proposed as a candidate for a synthetic genome project owing to its ease of propagation and rapidly growing toolbox for genetic engineering (Pampuch, Walker and Karas, 2022).

The preceding studies demonstrated that the chloroplast genomes of *Zea mays* (36.80% G+C) and *Chlamydomonas reinhardtii* (34.57% G+C) could be stably maintained in yeast and *E. coli*; however, the stability and maintenance of the *P. tricornutum* chloroplast genome (32.15% G+C) has yet to be explored and could pose unexpected challenges during cloning. Here, we generated a PCR-based and pre-cloned assembly strategy that demonstrated 90 to 100% efficiency when screening as few as ten yeast colonies and three *E. coli* colonies following whole-genome assembly and transformation, respectively.

## 2. RESULTS

### 2.1. Cloning of the P. tricornutum chloroplast genome

The PCR-based approach was first attempted as this enabled us to rapidly test various assembly setups by simply designing different sets of primers (Fig. 1, top panel). This resulted in an efficient assembly design that could be extrapolated to build the pre-cloned approach, which requires more time and energy to initially establish but permits greater downstream flexibility once generated (Fig. 1, bottom panel). Furthermore, the plasmids generated can be readily shared with other research facilities seeking to clone parts of or the whole *P. tricornutum* chloroplast genome.

**Figure 1.**
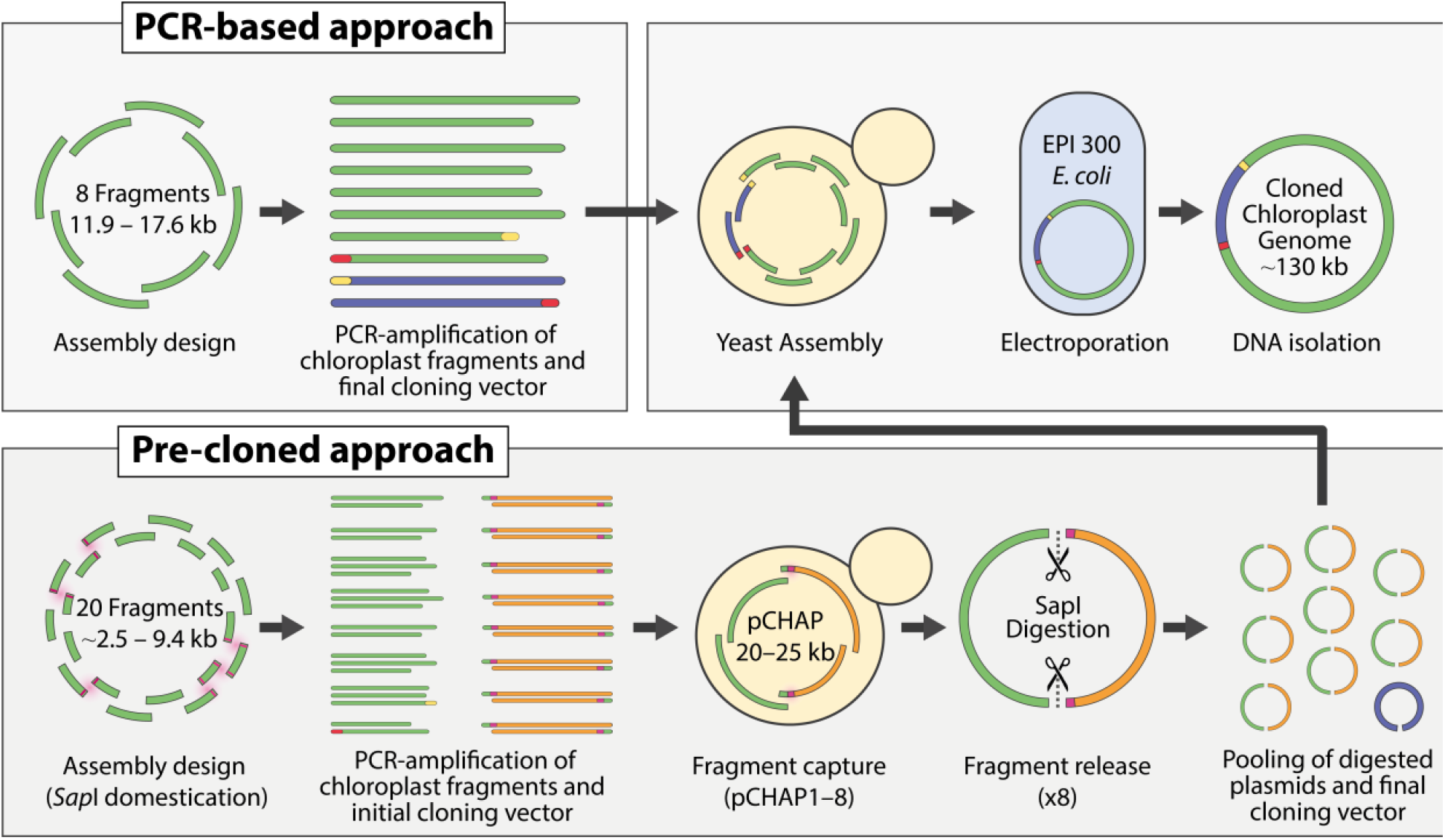
The two strategies for assembling the 117 kb chloroplast genome. The approaches differ in how the chloroplast fragments for whole-genome assembly are obtained. Endogenous and engineered *Sap*I restriction sites (magenta) are highlighted in the pre-cloned approach.

#### 2.1.2 Design strategy for partitioning the genome

It was first necessary to partition the 117 kb chloroplast genome into fragments that could be readily PCR-amplified. For this, three factors were considered while designing the constructs: (i) the inverted repeat regions (IRa and IRb, 6912 bp) should be flanked on either end by at least 1000 bp of non-repetitive DNA to avoid unwanted homologous recombination when attempting assembly of the whole chloroplast genome, (ii) fragment termini should overlap by at least 40 bp, but ideally within the range of 100 to 300 bp where possible, and (iii) the final cloning vector should be inserted into a non-coding region of the genome to reduce the possibility of interfering with endogenous gene expression in the final construct. Other design elements were considered depending on the objectives of the assembly strategy, as described in the proceeding sections.

#### 2.1.3 PCR-based assembly of the genome

We first partitioned the 117 kb chloroplast genome into fragments that could be easily obtained through PCR amplification. The fragments ranged in size from 11.9 to 17.6 kb and overlapped by 100 to 300 bp at all junctions except for the site where the cloning vector would integrate (Fig. 2A and B). Here, the respective fragments and cloning vector were amplified using primers that would generate 80 bp of overlapping homology at their termini (Table S1). The overlaps between the eight fragments and cloning vector were long enough to facilitate homologous recombination in *S. cerevisiae*.

**Figure 2.**
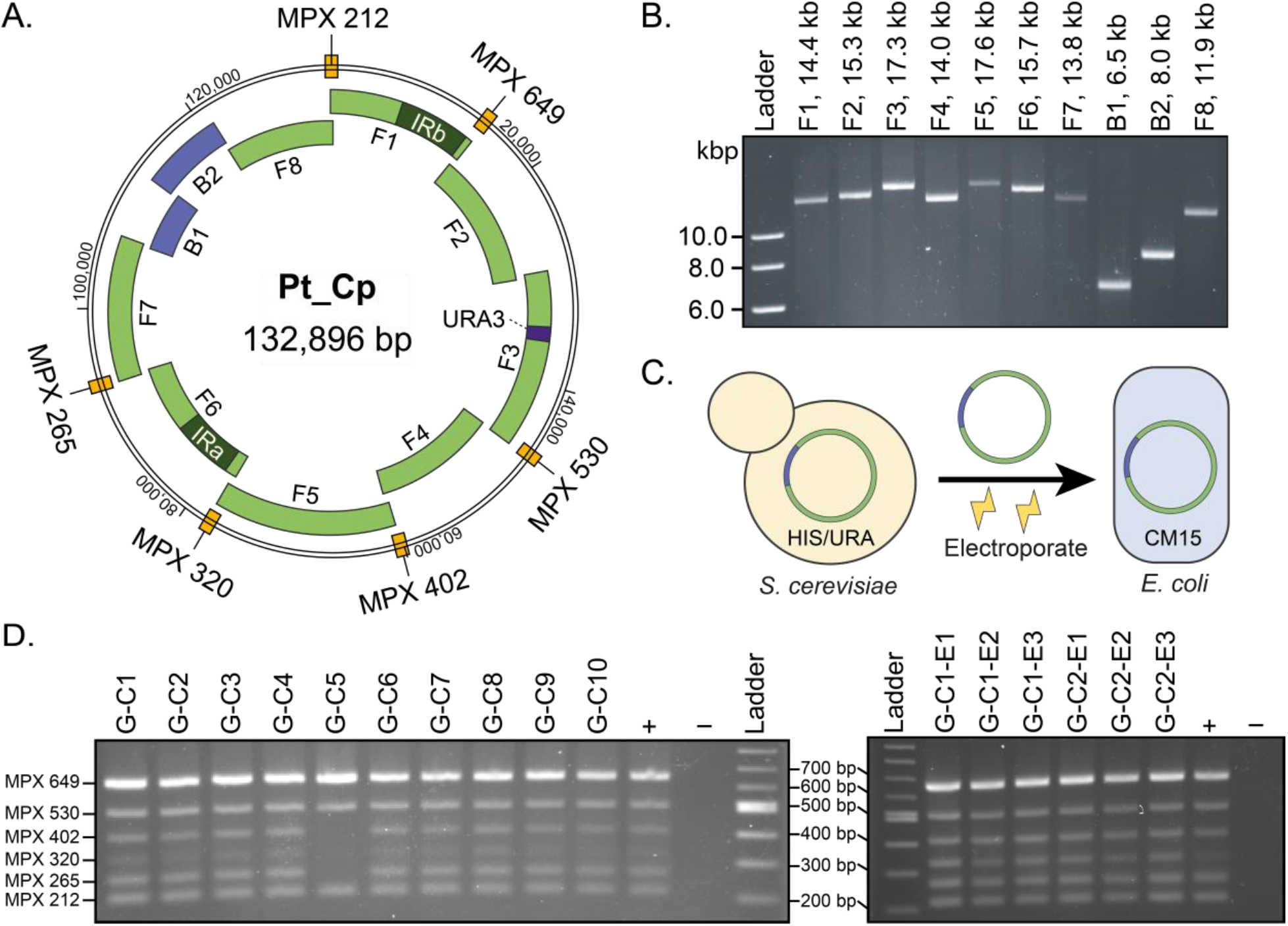
The assembly design for the PCR-based cloning method. **A**. The ten-fragment assembly includes the cloning vector (B1 and B2), which was split at the *HIS3* marker, and the eight chloroplast fragments. Fragment three was pre-cloned to have a second yeast selection marker, *URA3*, in a non-coding region of the genome. MPX amplicons were designed to span six of the eight junctions between chloroplast fragments. **B**. The PCR-amplified fragments used for whole-genome assembly. **C**. Plasmids from candidate *S. cerevisiae* colonies were electroporated into electrocompetent *E. coli*. **D**. Screening of ten yeast colonies following assembly of the whole genome. All but colony G-C5 screened positively. Plasmids from yeast colonies G-C1 and G-C2 were electroporated into *E. coli*, with all six transformants screening positively. Amplicons were visualized on a 1% agarose gel stained with ethidium bromide.

We tested various integration sites and found that the non-coding region between *ycf88* and *rpl22*, which maps to the junction between chloroplast fragments seven and eight, was the most easily amplifiable. After a few failed attempts, we also integrated the yeast selective marker *URA3* into one of the chloroplast fragments to increase the probability of correct assembly of the whole chloroplast genome. The *URA3* marker was amplified using 80 bp primers that would add 40 bp of homology at either termini to a non-coding region in chloroplast fragment three. The third fragment was resultantly split into three smaller fragments, annotated as fragments 3A, 3B, and 3C (Table S1). A 13-fragment assembly was performed in *S. cerevisiae* and demonstrated that the whole chloroplast genome could be recovered when using two yeast selective markers interspersed throughout the genome. To reduce the number of fragments needed for assembly, thereby increasing efficiency, the *URA3* marker was pre-cloned into chloroplast fragment three, generating the plasmid pCHAP3_URA (Fig. S1A and B). From this plasmid, the third fragment containing the integrated marker could be amplified as an intact piece via PCR and used in a 10-fragment assembly to reconstitute the whole genome.

The 10-fragment PCR-based assembly demonstrated that nine out of ten of *S. cerevisiae* colonies screened positively, as per the multiplex PCR, when selected at random (Fig. 2D). When we electroporated the plasmids from two candidate *S. cerevisiae* colonies into *E. coli* (Fig. 2C), all the analyzed transformants screened positively when as few as three transformants were screened per transformation (Fig. 2D). Plasmid DNA was isolated from *E. coli* colonies G-C1-E1 and G-C2-E1 and sequenced, revealing that the whole chloroplast genome had been successfully captured through this method.

#### 2.1.4 Pre-cloned approach for assembling the genome

The design strategy identified through the PCR-based approach was used to inform the pre-cloned approach. Here, we first PCR amplified the chloroplast genome as 20 overlapping fragments ranging from 2.5 to 9.4 kb in length (Table S1). Fragment termini were strategically placed at the six endogenous *Sap*I recognition sites so that primers could be used to introduce silent mutations, thereby domesticating the chloroplast genome for *Sap*I (Fig. S2A). Pairs or triads of fragments were then assembled with the cloning vector pSAP1, which had been previously domesticated for *Sap*I using the same process. The plasmids were designed such that the chloroplast fragments could be scarlessly released from their respective plasmids upon digestion with *Sap*I (Fig. S2B). This ultimately generated eight unique plasmids, pCHAP1 to 8, each harbouring one of the eight overlapping chloroplast fragments identified through the PCR-based approach.

We also domesticated the cloning vector pDMI_2, generating the plasmid pDMI_Sap. This plasmid was then assembled to have a *Sap*I recognition site flanked by 28 bp homologies to chloroplast fragments seven and eight, generating pDMI_7/8 (Fig. 3A). When digested with *Sap*I, pDMI_7/8 becomes linearized, exposing the 28 bp homologies at its termini. Plasmids pCHAP7 and pCHAP8 were designed to contain 20 bp homologies to the respective pDMI_7/8 termini, making it such that in total, the fragments shared 48 bp of overlap at the respective termini. The eight pCHAP plasmids and cloning vector pDMI_7/8 were digested, both individually and in a one-pot reaction (Fig. 3B), before transformation into *S. cerevisiae* for assembly of the whole genome.

**Figure 3.**
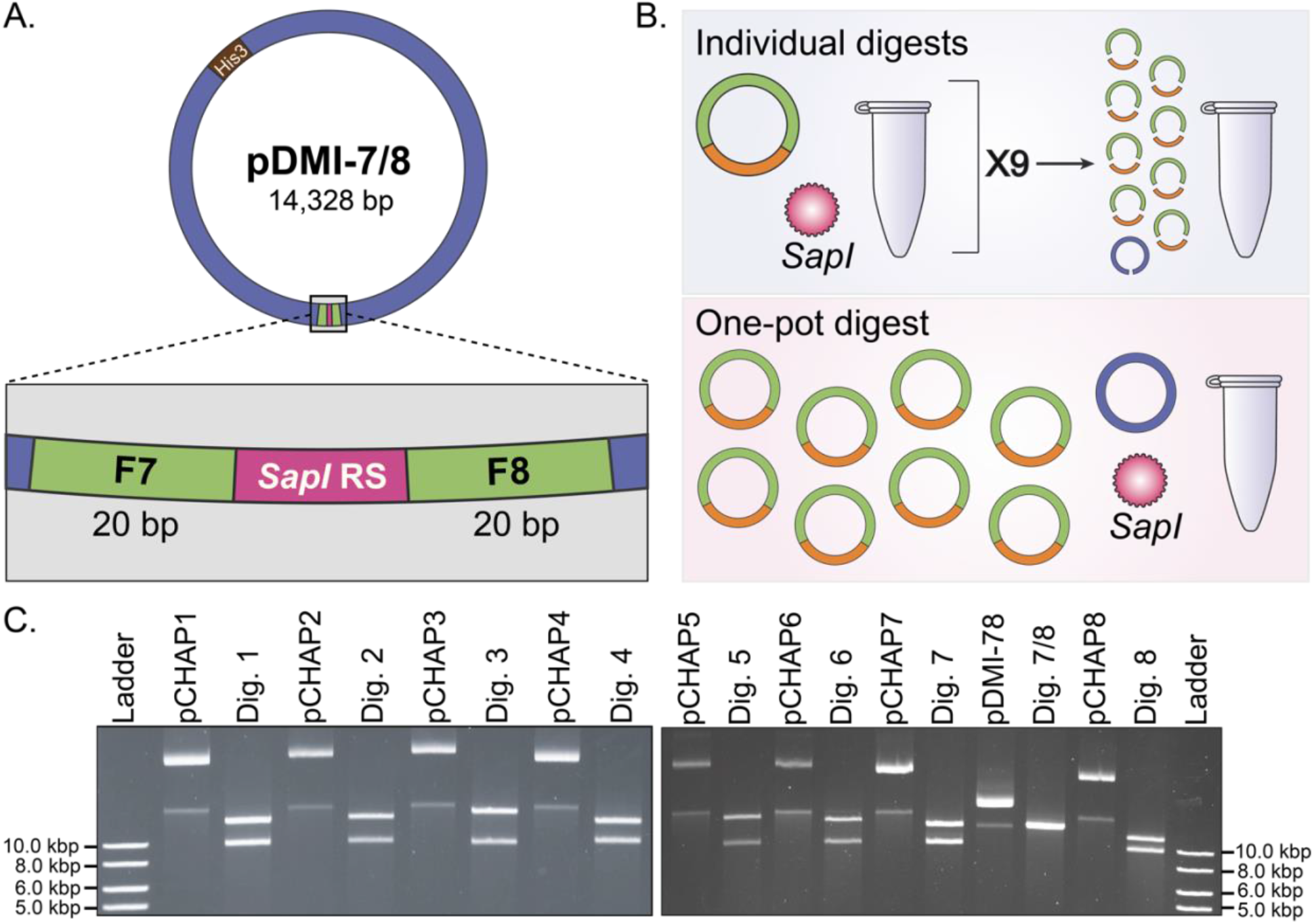
Plasmid-based assembly of the *P. tricornutum* chloroplast genome. **A**. pDMI_7/8 was designed to have a single *Sap*I recognition site (RS) flanked by 20 bp homologies to chloroplast fragments seven and eight. When digested with *Sap*I, pDMI_7/8 is linearized, exposing the homologies necessary for assembly. **B**. Two assembly mechanisms were possible – one in which all the plasmids are individually digested and validated ahead of assembly, and another in which all the plasmids are digested simultaneously in a one-pot reaction. The latter is more time-effective but cannot be assayed to ensure digestion is complete. **C**. Plasmids pCHAP_1 to pCHAP_8 and pDMI-7/8 were digested until completion via *Sap*I, liberating the individual chloroplast fragments from the cloning vector pSAP1 as well as linearizing the final cloning vector.

The individual reactions were assayed on a gel to ensure digestion was complete (Fig. 3C).

The pre-cloned assembly approach demonstrated that nine out of ten (individual digests, Fig. S3A) and ten out of ten (one-pot digest, Fig. S3B) *S. cerevisiae* colonies screened positively, as per the multiplex PCR, when selected at random. Plasmids from two candidate *S. cerevisiae* colonies per assembly were electroporated into *E. coli*, whereupon all the analyzed transformants screened positively (Fig. S3). Plasmid DNA from *E. coli* colonies S-C2-E1 and C-C1-E1 was isolated and then sequenced, revealing that the whole chloroplast genome had been successfully reconstituted using both setups for the pre-cloned assembly method.

### 2.2 Maintenance of P. tricornutum chloroplast plasmids in E. coli

A growth assay was performed to determine if the cloned chloroplast genome poses a burden to EPI300 *E. coli*. Growth was measured in the presence and absence of arabinose, which induces expression of *trfA* via the pBAD promoter when present, leading to high-copy number replication of the genome. The growth rate of strains containing the chloroplast genome was measured against strains harbouring pSAP1 and pDMI_7/8, which were the cloning vectors used for initial fragment capture and whole-genome cloning, respectively. Under non-induced conditions (i.e., no arabinose present), the strains demonstrate a similar growth rate (Fig. 4A); however, when plasmid replication is induced via arabinose, the strains harbouring the genome grow substantially slower and to a lower density than the strains harbouring the comparatively small cloning vectors (Fig. 4B).

**Figure 4.**
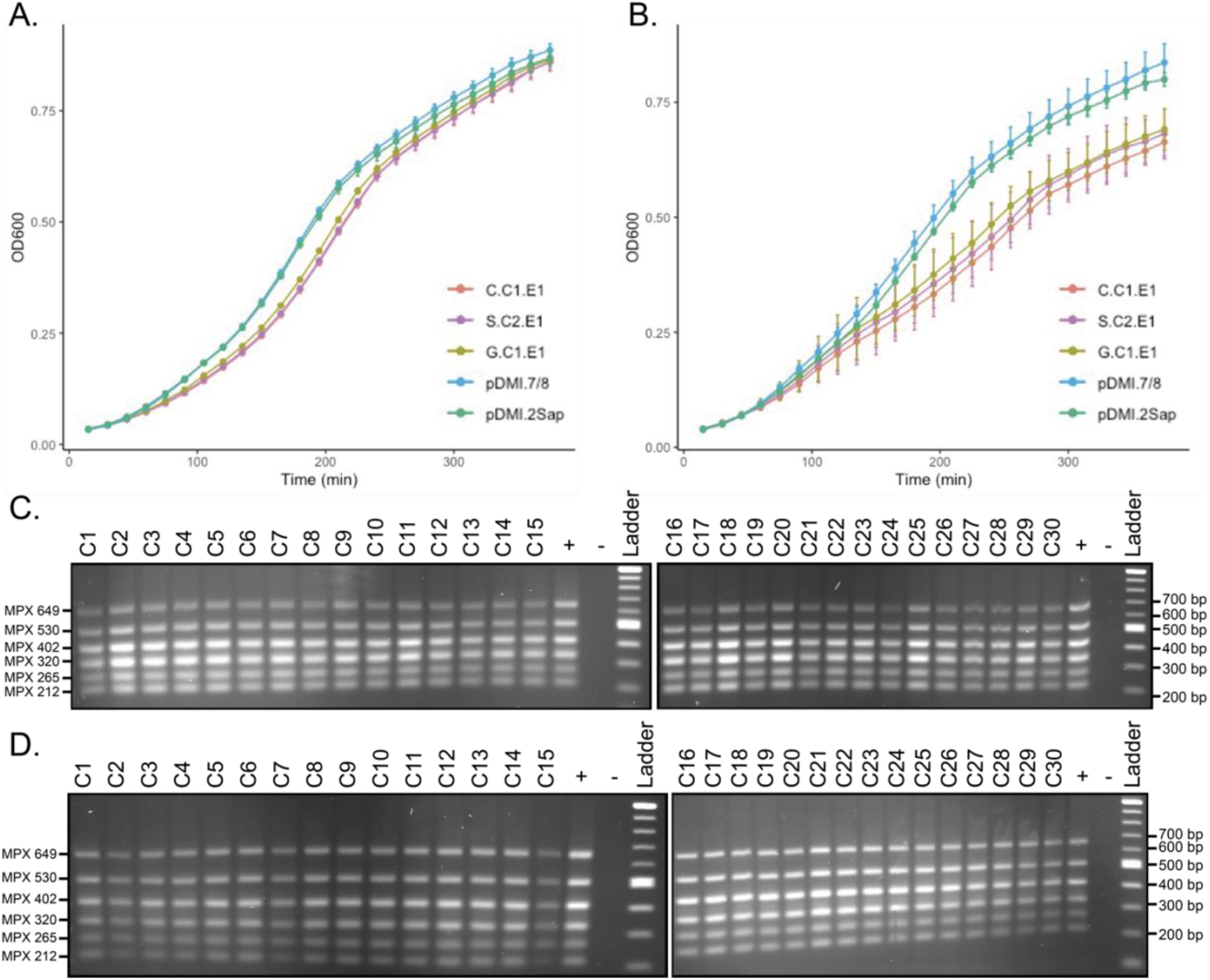
Analysis of the burden and stability of the cloned *P. tricornutum* chloroplast genome in *E. coli*. **A**. Under standard conditions (i.e., LB broth supplemented with CM15, 37ºC), *E. coli* strains harbouring the chloroplast genome do not exhibit a growth deficit when compared to strains harbouring smaller plasmids (∼14 kb). **B**. When *E. coli* are induced to high-copy-number plasmid replication, strains harbouring the chloroplast genome demonstrate a substantial growth burden. **C**. MPX screening of 30 colonies isolated from the *E. coli* strain harbouring G-C1-E1 at time-point zero. All colonies demonstrate the expected banding pattern. **D**. MPX screening of 30 colonies isolated from the *E. coli* strain harbouring G-C1-E1 at time-point 60 (i.e., 60 generations). All colonies demonstrate the expected banding pattern. Amplicons were visualized on a 1% agarose gel stained with ethidium bromide.

Following this, a stability assay was conducted to see if the cloned genome can be stably maintained over the course of several bacterial generations. Thirty clones harbouring the first-sequenced chloroplast genome (i.e., G-C1-E1) were grown for approximately sixty generations. DNA was isolated and screened from the clones prior to (Fig. 4C) and directly after (Fig. 4D) subculturing for this length of time. The MPX screen suggested that none of the clones had any discernable rearrangements or large-scale deletions within the plasmid, at least in the regions spanning six of the eight possible junctions between chloroplast fragments. The first three clones had their plasmids sequenced, demonstrating that the chloroplast genome had been maintained without any major rearrangements or deletions, as the MPX screen had suggested.

## 3. DISCUSSION

We have developed a strategy for whole genome assembly that results in 90 to 100% efficiency when selecting as few as ten *S. cerevisiae* colonies and three *E. coli* colonies following assembly and electroporation, respectively. Our strategy was first optimized using a PCR-based method, which allowed for the rapid testing of various assembly designs, before being adapted into the pre-cloned approach. The latter assembly strategy enables greater downstream flexibility as the pre-cloned fragments of the chloroplast genome can be further manipulated within *S. cerevisiae* (e.g., TREC-assisted gene knock-in; Chandran *et al*., 2014) or *E. coli* (e.g., lambda-red recombineering; Sharan *et al*., 2009) before attempting whole-genome assembly.

Initial attempts to assemble the genome in *S. cerevisiae* failed. This could have been due to recombination occurring between the inverted repeats (i.e., IRa, IRb) endogenously present within the chloroplast genome, or simply because of the size and number of fragments involved, which increased the complexity of the reaction. To generate greater selection pressure for the correct assembly, we incorporated a second yeast selective marker for the uracil biosynthetic pathway (i.e., *URA3*) into a non-coding region of the long single copy (LSC). The second marker was positioned approximately 57.7 kb away from the cloning vector harbouring a histidine biosynthetic marker (i.e., *HIS3*) in the small single copy (SSC). The LSC and SSC are delineated by the inverse repeat regions; by positioning yeast selection markers on either side of this delineation, we aimed to encourage correct assembly of the SSC and LSC in *S. cerevisiae* when selecting on -HIS/URA media. This design proved successful at reconstituting the whole genome.

For the second assembly approach, fragments of the chloroplast genome were domesticated for endogenous *Sap*I sites and pre-cloned into plasmids. The initial design for the whole genome was entirely domesticated for *Sap*I, which required domesticating the cloning vector pDMI_7/8 and the integrated *URA3* marker as well. However, the endogenous *Sap*I restriction site in *URA3* poses an interesting opportunity for engineering, and so a new design was created and assembled. Pre-cloned and PCR-amplified fragments were combined to generate a cloned chloroplast genome that harboured a single *Sap*I site in the *URA3* marker. This restriction site can be used to integrate transgenic cassettes into the chloroplast genome through golden gate cloning or yeast assembly of the linearized plasmid and a transgenic fragment containing the appropriate overlaps (Fig. S2C). The resulting transformed yeast can be selected for on

-HIS media supplemented with 5-fluorotic acid (5-FOA). If *URA3* is still intact following assembly, 5-FOA will be metabolized into fluorouracil, thereby killing yeast with unsuccessfully assembled plasmids. This provides a simple and efficient method for rapidly integrating any transgenic cassette into the cloned chloroplast genome.

The cloned genomes were able to be stably maintained in *E. coli* and only perturbed growth rates when induced to high-copy number replication via supplementation of the media with arabinose. The cloning of partial-or whole-prokaryotic-derived genomes can pose issues within *E. coli* due to the unwanted expression of genes, which can pose energetic costs or even toxicity to the cell, or through competition for DNA replication and partitioning machinery between endogenous and introduced genetic material (Godiska *et al*., 2005). Another stability concern was that the *P. tricornutum* chloroplast genome has low G+C content (32.5%) relative to the *E. coli* genome (50.8%). Plasmids that are A+T rich can be challenging to clone in *E. coli* as these regions may spuriously form genetic elements like promoters, replication origins, open reading frames, and more (Godiska *et al*., 2005). In this instance, we believe that the decrease in growth rate under the induced growing condition is likely due to the increased energy-demand associated with replicating several copies (e.g., tens to hundreds) of a large plasmid. The stability and replicability of the chloroplast genome under non-induced growing conditions presents a potential mechanism for delivering the cloned genome to *P. tricornutum* via bacterial conjugation (Karas *et al*., 2015). However, it is thought that conjugated plasmids are specifically targeted to the eukaryotic-cell nucleus during this process; further engineering efforts will be required to realize the potential for conjugation as a transformation method for organelle engineering. In the meantime, the chloroplast genome could be transformed into *P. tricornutum* through biolistic bombardment or PEG-mediated transformation, which have been used to introduce foreign DNA into the chloroplasts of other microalgal species.

Before attempting to deliver the chloroplast genome through conjugation or any other transformation method, it will be necessary to disrupt *RecA* and incorporate a chloroplast-specific selection marker into the cloned genome. The bacterial-derived recombinase *RecA* (PHATRDRAFT_54013) is expressed in the nuclear genome but localizes to the chloroplast after translation, where it mediates homologous recombination. Previous studies have demonstrated that disrupting this gene in a variety of distantly related photosynthetic eukaryotes disrupts homologous recombination within the chloroplast (Cerutti *et al*., 1995; Jeon *et al*., 2013). This will be necessary to prevent homologous recombination between the endogenous and cloned genomes, which posed challenges during transformation of the *Chlamydomonas* chloroplast genome (O’Neill *et al*., 2011). Many chloroplast-specific markers have been demonstrated in photosynthetic eukaryotes, but to-date, only the chloramphenicol resistance marker *cat* has been explored in *P. tricornutum* (Xie *et al*., 2014). It will be worthwhile to explore other resistance markers, like the commonly used streptomycin-resistance marker *aadA*, to expand the organelle engineering toolbox for *P. tricornutum*.

In summary, we developed two approaches for efficient assembly of the *P. tricornutum* chloroplast genome and a facile mechanism for integration of transgenic cassettes, enabling the rapid generation of designer genomes. The chloroplast genome was able to be stably maintained and propagated in *E. coli* when plasmid replication was maintained at a low-copy number (i.e., non-induced). To realize the full potential of this assembly pipeline, it will be necessary to develop methods for delivering, selecting for, and maintaining the cloned genome inside the recipient cell’s chloroplast. The design-build cycle described herein could be adapted to generate *P. tricornutum* chloroplast genomes of any biologically conceivable imagination (Fig. 5) or even extrapolated to clone the chloroplast genomes of other photosynthetic eukaryotes (e.g., *Thalassiosira pseudonana*).

**Figure 5.**
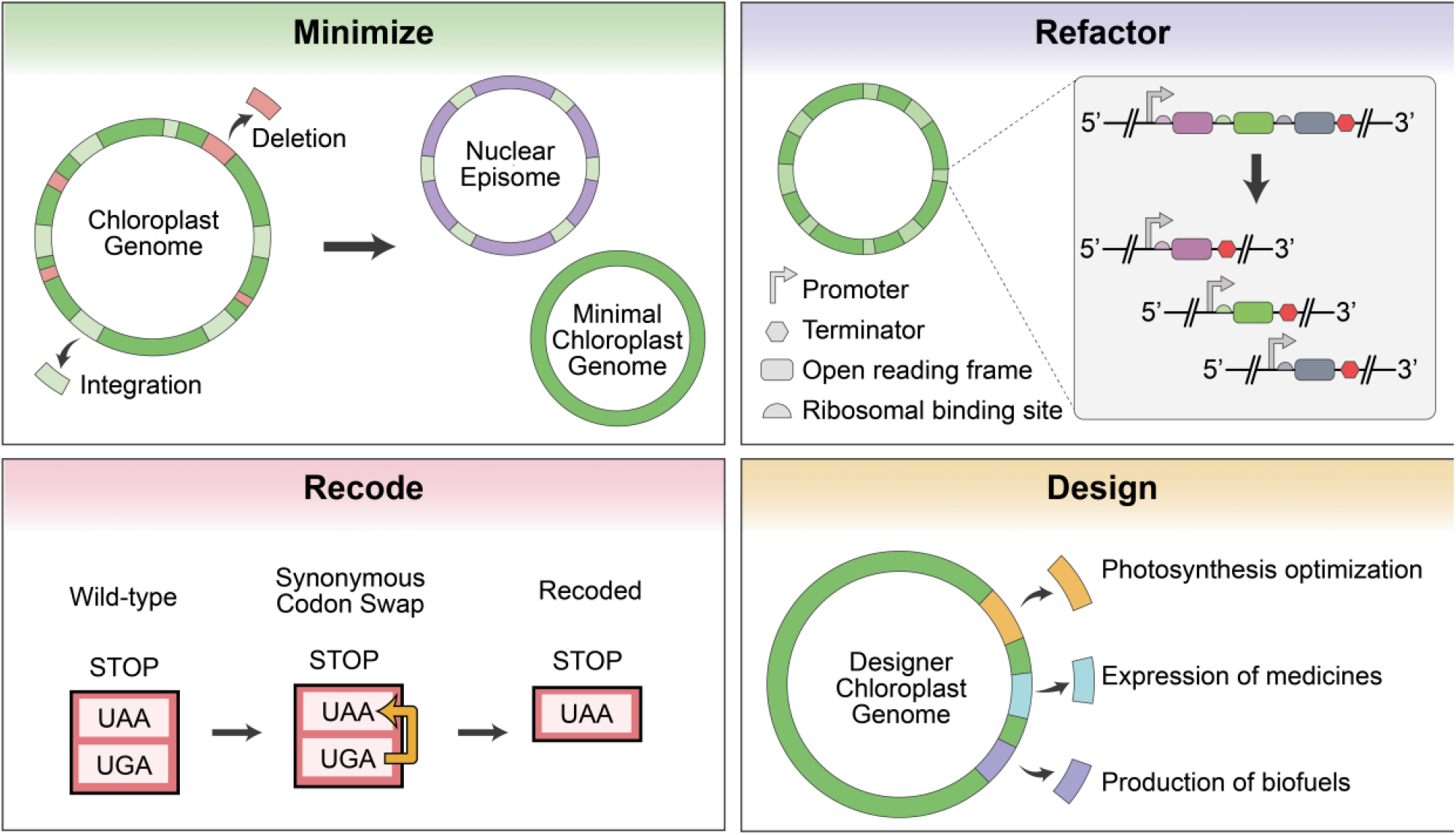
Potential applications for future designer chloroplast genomes (adapted from Fig. 6 of Esvelt and Wang (2013). Genome minimization can be achieved by deleting nonessential regions of the genome and integrating essential genes into an episome for nuclear expression. Tractability can be increased by refactoring operons so that every gene has a unique promoter and terminator. Recoding of the stop codon UGA to UAA could facilitate codon reassignment to a non-natural amino acid for biocontainment purposes. Taken altogether with the proposed methods, it will be possible to design the chloroplast genome for several purposes.

## AUTHOR CONTRIBUTIONS

EJLW oversaw and performed most of the experimental work, created the figures, and wrote the manuscript. MP and NC aided with experiments while under the mentorship of EJLW and provided edits to the manuscript. As well, MP created the growth assay graphs and performed statistical analyses. RC performed preliminary unpublished work that would serve as the starting point for designs presented within this manuscript. BK conceived the project, assisted in the design, and helped prepare the manuscript.

## ACKNOWLEDGEMENTS

Daniel Giguere provided high-quality template DNA for *P. tricornutum* that was used to troubleshoot the initial PCR reactions. Flow Genomics and Plasmidsaurus sequenced the cloned genomes. This work was supported by Natural Sciences and Engineering Research Council of Canada (RGPIN-2018-06172) awarded to B.J.K.

